# Multi-COBRA hemagglutinin formulated with cGAMP microparticles elicit protective immune responses against influenza viruses

**DOI:** 10.1101/2024.02.27.582355

**Authors:** Xiaojian Zhang, Hua Shi, Dylan A. Hendy, Eric M. Bachelder, Kristy M. Ainslie, Ted M. Ross

## Abstract

Influenza viruses cause a common respiratory disease known as influenza. In humans, seasonal influenza viruses can lead to epidemics, with avian influenza viruses of particular concern because they can infect multiple species and lead to unpredictable and severe disease. Therefore, there is an urgent need for a universal influenza vaccine that provides protection against seasonal and pre-pandemic influenza virus strains. The cyclic GMP-AMP (cGAMP) is a promising adjuvant for subunit vaccines that promotes type I interferons production through the stimulator of interferon genes (STING) pathway. The encapsulation of cGAMP in acetalated dextran (Ace-DEX) microparticles (MPs) enhances its intracellular delivery. In this study, the Computationally Optimized Broadly Reactive Antigen (COBRA) methodology was used to generate H1, H3, and H5 vaccine candidates. Monovalent and multivalent COBRA HA vaccines formulated with cGAMP Ace-DEX MPs were evaluated in a mouse model for antibody responses and protection against viral challenge. Serological analysis showed that cGAMP MPs adjuvanted monovalent and multivalent COBRA vaccines elicited robust antigen-specific antibody responses after a prime-boost vaccination and antibody titers were further enhanced after second boost. Compared to COBRA vaccine groups with no adjuvant or blank MPs, the cGAMP MPs enhanced HAI antibody responses against COBRA vaccination. The HAI antibody titers were not significantly different between cGAMP MPs adjuvanted monovalent and multivalent COBRA vaccine groups for most of the viruses tested in panels. The cGAMP MPs adjuvanted COBRA vaccines groups had higher antigen-specific IgG2a binding titers than the COBRA vaccine groups with no adjuvant or blank MPs. The COBRA vaccines formulated with cGAMP MPs mitigated disease caused by influenza viral challenge and decreased pulmonary viral titers in mice. Therefore, the formulation of COBRA vaccines plus cGAMP MPs is a promising universal influenza vaccine that elicits protective immune responses against human seasonal and pre-pandemic strains.

## Introduction

The World Health Organization (WHO) estimates that annually influenza-induced respiratory disease results in ∼650,000 deaths worldwide (1, 2). Due to its segmented, negative-sense viral RNA genome, influenza virus evolves rapidly into multiple variants each year (3). Immune responses directed against the viral proteins hemagglutinin (HA) and neuraminidase (NA) on the surface of the virion provide protection against virus infection (4–6). However, the HA and NA proteins are highly variable proteins with 18 HA and 11 NA subtypes. Seasonal influenza type A (H1N1 and H3N2) and type B (Yamagata and Victoria lineages) viruses cause epidemics in humans, while avian influenza viruses (*e.g.* H5Nx and H7N9) can cross species and cause severe disease in humans due to the avian receptors (alpha-2,3 sialidase receptors) present in the lower respiratory tract of humans.

Annual vaccination is currently the primary method for controlling the spread and impact of influenza virus-induced disease. To match the antigenicity of current circulating viral variants, the strains used in the annual seasonal influenza vaccine formulation are updated based on global surveillance. Even with annual reformulation, the effectiveness of influenza vaccines is between 10-60% (https://www.cdc.gov/flu/vaccines-work/past-seasons-estimates.html). Thus, there is an urgent need for an improved next-generation influenza vaccine that increases broadly protective immune responses against influenza viral infection to not only better protect against a potential pandemic, but also reduces the annual manufacturing burden of new seasonal influenza vaccine. As outlined by Fauci et al. in 2018 (7), a universal influenza vaccine should have the coverage of vaccine protection against all influenza A viruses (with or without influenza B viruses). The design strategy of many universal influenza vaccines is to direct the immune response to conserved protective epitopes on HA, NA, NP, and M2 proteins (8–13). The COBRA design is a multi-layer consensus building approach (9, 14–16), that utilizes HA protein sequences from historical and circulating influenza viral isolates to generate a COBRA HA protein. COBRA-based vaccines direct the immune response towards epitopes on the globular head domain of both HA and NA glycoproteins. Our previous studies have shown that COBRA HA vaccines are superior wild type HA vaccines in coverage of protection (10, 14–23), indicating that the COBRA methodology is an effective vaccine design strategy for universal influenza vaccine development. Further, COBRA HA influenza antigens are highly compatible with multiple vaccine platforms. With reverse-genetic technology, sequences of COBRA antigens can be used to generate the COBRA-based reassortant viruses that then can be used to develop inactivated and live-attenuated influenza vaccines, which commercially available platforms used to make influenza virus vaccines today. In addition, COBRA HA antigens can also take the advantage of mRNA vaccine platform, which has achieved great success in the combat against COVID-19 pandemic (24).

The current quadrivalent influenza vaccine is a split inactivated influenza virus (IIV; e.g. Fluzone), recombinant HA protein (Flublok) or live-attenuated influenza virus (LAIV; FluMist) vaccine that contains one of each H1N1 and H3N2 influenza A virus and two influenza B viruses (Yamagata and Victoria lineages). All these vaccines induce strong immune responses against homologous influenza viruses, but are less effective in controlling influenza virus infection by antigenic drifted strains or pre-pandemic strains. The IIV vaccine also has high-dose (HD) and adjuvanted (MF59) versions that are licensed for people 65 years and older. Often people in this age group may be immunocompromised or have weakened immune systems and therefore additional stimulation, such as an adjuvant, is needed to enhance the immune responses during vaccination.

The use of adjuvants that stimulate the innate immune system enhance influenza vaccine elicited immune responses (25). Often vaccine adjuvants activate innate immune cells (*e.g*. dendric cells) through pattern recognition receptors (PRRs) that recognize pathogen- or damage-associated molecular patterns (PAMPs or DAMPs) (25). PRRs include toll-like receptors (TLRs), cytosolic receptors, and C-type lectin receptors (reviewed in (25)). One such cytosolic receptor is the stimulator of interferon genes (STING) PRR that activates in response to cyclic dinucleotides (CDNs). Activation of the STING pathway stimulates interferon regulatory factor 3 (IRF3) and nuclear factor-κB (NF-κB), which results in the transcription of type I interferons (IFNs) and other pro-inflammatory cytokines to modulate antigen presentation and immune responses. There are several CDN agonists of STING. First, 2’ 3’-cyclic guanosine monophosphate-adenosine monophosphate (2’ 3’-cGAMP), which is synthesized by the enzyme cyclic guanosine monophosphate-adenosine monophosphate synthase (cGAS). Second, 3’ 3’ -cGAMP, is produced by numerous bacterial pathogens and activates STING. Finally, other CDN STING agonists include cyclic dimeric guanosine monophosphate (c-di-GMP) and cyclic dimeric adenosine monophosphate (c-di-AMP). This range of STING agonists highlights their potential as initiators for the innate immune response, which is crucial for the effectiveness of subunit vaccines (26).

However, because of their negative charge and hydrophilicity, extracellular CDNs do not penetrate well through the cellular membrane to reach their cytosolic STING receptor, even when high doses of soluble CDNs are used (27). Compared to soluble form, the CDNs encapsulated in liposomes and polymeric particles have been reported to enhance bioactivity (28–32). To enhance CDN delivery and long-term stability, cGAMP was formulated into acetalated dextran (Ace-DEX) polymeric microparticles (MPs) by electrohydrodynamic spraying (electrospray) (33–35). Ace-DEX is synthesized from FDA-approved water-soluble dextran by converting the pendant hydroxyl groups into acetal groups, forming a hydrophobic polymer that is biocompatible, biodegradable, and stable at elevated temperatures (33). When formed into MPs through electrospray, high adjuvant encapsulation efficiency can be achieved that display dose sparing of the encapsulated adjuvant (34–37). Ace-DEX MPs are sized (∼1-2 microns) to passively target phagocytic cells (e.g. dendritic cells) and the polymer’s acid sensitivity facilitates enhanced release of payload in the acidic environment of the phagolysosome once phagocytosed (33, 38, 39).

cGAMP encapsulated in Ace-DEX MPs (cGAMP MPs) induced enhanced type-I interferon responses *in vitro* and *in vivo*, balanced Th1- and Th2-associated responses, and expanded germinal center B cells and memory T cells, as compared to soluble cGAMP (34). Further, a cGAMP MP adjuvanted subunit HA vaccine induced protective immunity against a lethal influenza challenge in both mice and ferrets, with no observed toxicity in vaccinated animals (34, 40). However, only a monovalent wild-type or COBRA H1 or H3 HA subunit vaccines were tested with Ace-DEX cGAMP MPs (34, 40–45). The ability of cGAMP MPs to generate a broadly reactive vaccine response with multiple universal influenza vaccine antigens has yet to be studied. In this study, the COBRA-based vaccines (H1, H3, and H5) were designed to overcome the antigenicity diversity of influenza viruses and their effectiveness as a subunit vaccine was evaluated with and without cGAMP MPs in a mouse model.

## Materials and Methods

### Hemagglutinin, cGAMP microparticles and vaccination

The HA antigens J4 (23), Y1 (16), and IAN5 (46) were designed using the next-generation COBRA methodology (15). Briefly, full length wild-type influenza A(H1N1), A(H3N2) and A(H5Nx) HA protein amino acid sequences were downloaded from public online databases. The J4 (H3 COBRA) HA was derived from sequences isolated between 2013 - 2016, Y1 (H1 COBRA) HA was derived from sequences isolated between 2013 - 2015, and IAN5 (H5 COBRA) HA was derived from sequences collected between 2011 - 2015 (Fig. 1A).

**Figure 1.**
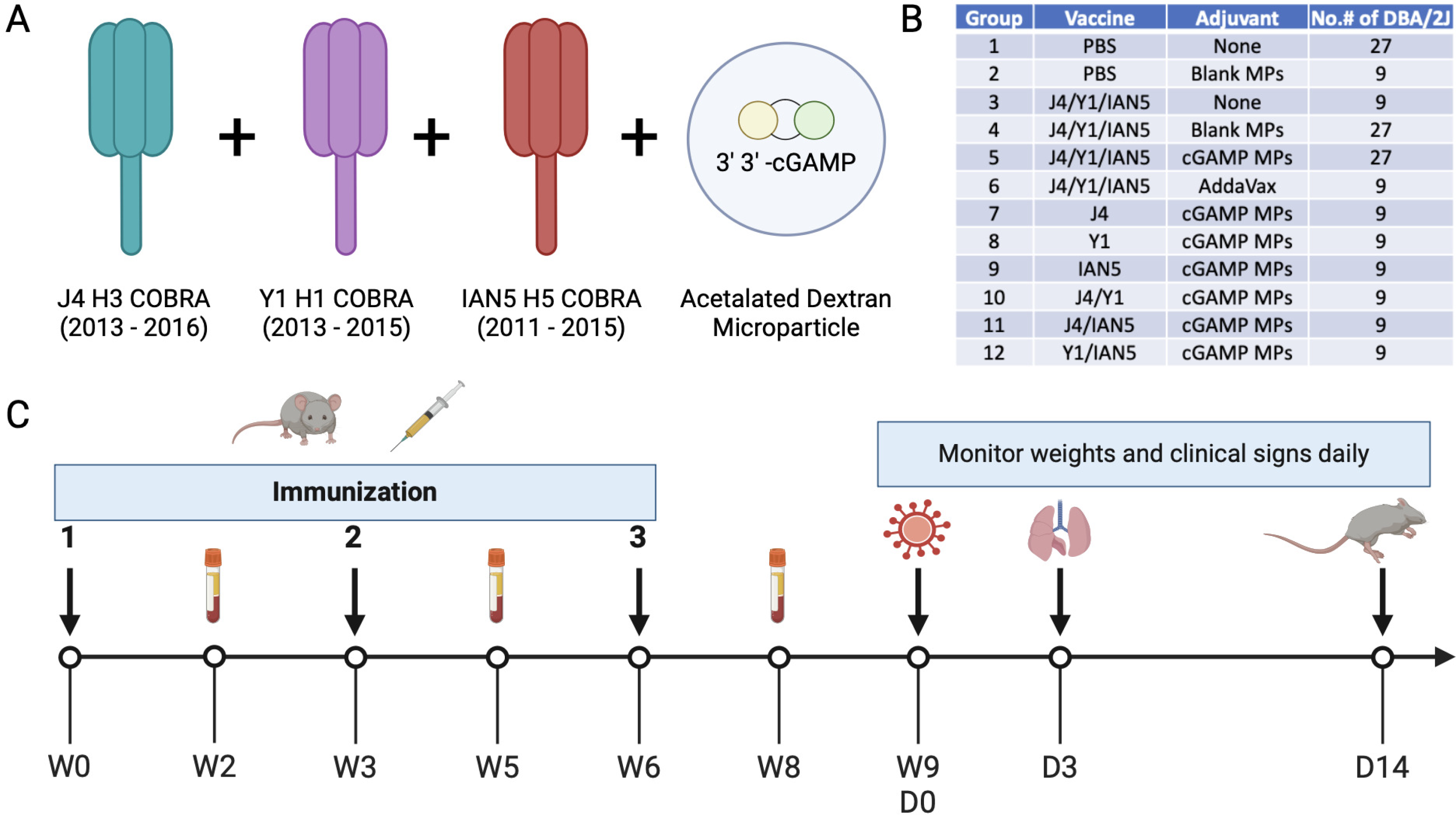
Experimental design. (A) Vaccine formulation. Antigens J4, Y1, and IAN5 COBRAs were co-administrated with cGAMP encapsulated in Ace-DEX microparticles (MPs). (B) Experimental groups. (C) Schedule for vaccination and challenge. Female DBA/2J mice were vaccinated three times, three weeks apart, with the same vaccine formulation. Sera samples were collected after each vaccination for analysis. The mice were challenged with either H1N1, H5N6 or H3N2 influenza viruses at 3 weeks after final boost. Lung samples were collected on day 3 post-infection for pulmonary viral loads. Over the course of infection, weights and clinical signs were monitored for 14 days.

Soluble HA proteins were obtained by transfecting truncated HA genes that were cloned into the pcDNA3.1+ plasmid into HEK293T suspension cells as previously described (47). The truncated HA genes were generated by replacing the transmembrane domain with a T4 fold-on domain, an Avitag, and a 6× His-tag for purification. The concentration of the soluble HA proteins was determined by conventional bicinchoninic acid assay (BCA) according to the manufacture’s instruction.

Ace-DEX was synthesized according to Kauffman *et al.* using dextran from *Leuconostoc mesenteroides* (molecular weight = 70 kDa; Sigma, St. Louis, MO, USA) (33). Polymer was formed into MPs via electrospray to form blank MPs or encapsulate cGAMP (cGAMP MPs) as previously reported (44). cGAMP loading in MPs was analyzed via HPLC and determined to be 10.86 µg cGAMP/mg total. Endotoxin contamination was also evaluated in the MPs via the Pierce Chromogenic Endotoxin Quant Kit and found to be below the limit of detection (<0.1 EU/mL). The morphology of the MPs was analyzed via scanning electron microscopy (Hitachi s-4300 Cold Field Emission SEM; Ibaraki, Japan; UNC CHANL) (SFig. 1A-B).

DBA/2J mice (females, 6-8 weeks old) were purchased from The Jackson Laboratory (Bar Harbor, ME, USA) and housed in microisolator units with the free access to food and water. All animals were cared for under the USDA guidelines for laboratory animals, and all procedures were approved by the University of Georgia Institutional Animal Care and Use Committee (IACUC) (no. A2021 06-016).

Influenza naïve DBA/2J mice (9/group) were vaccinated with a mixture of COBRA rHA (1 μg HA per antigen) and blank or cGAMP MPs (1 μg cGAMP per mouse) in a total volume of 50 μL PBS delivered intramuscularly in the thigh. COBRA rHA (1 μg per antigen) with no adjuvant served as an antigen only vaccination. PBS with no adjuvant served as a mock vaccination. The AddaVax squalene-based oil-in-water adjuvant (InvivoGen, San Diego, CA, USA) was used as an adjuvant control. Mice were vaccinated at week 0 and boosted with the same vaccine formulation at the same dose at weeks 3 and 6. Blood samples were collected from mice via sub-mandibular bleeds 2 weeks after each vaccination in 1.5-ml microcentrifuge tubes. The samples were incubated at RT for 30 min and then centrifuged at 6,000 rpm for 5 min. Serum samples were collected and stored at -20°C until further analysis.

### Viruses and viral challenge

The H1, H3, and H5 subtype viruses were obtained from either the Influenza Reagents Resource (IRR), BEI Resources, or the Centers for Disease Control and Prevention (CDC). Viruses were propagated once in either embryonated chicken eggs or Madin-Darby canine kidney (MDCK) cells (Sigma). To verify the presence of newly propagated viruses, hemagglutination titer of the frozen aliquots was determined with either 0.8% turkey red blood cells (TRBCs) for H1 subtype viruses, 0.75% guinea pig red blood cells (GPRBCs) (Lampire Biologicals, Pipersville, PA, USA) for H3 subtype viruses in the presence of 20nM Oseltamivir, or 1% horse red blood cells (HoRBCs) for H5 subtype viruses. Virus lots were aliquoted for single-use applications and stored at -80°C. The H1 virus panel includes: A/Victoria/2570/2019 (Vic/19, H1N1), A/Guangdong/SWL1536/2019 (GD/19, H1N1), A/Brisbane/02/2018 (Bris/18, H1N1), A/California/07/2009 (Cal/09, H1N1), A/Brisbane/59/2007 (Bris/07, H1N1). The H3 virus panel includes: A/Darwin/9/2021(Dar/21, H3N2), A/Tasmania/503/2020 (Tas/20, H3N2), A/Hong Kong/2671/2019 (HK/19, H3N2), A/South Australia/34/2019 (SA/19, H3N2), A/Kansas/14/2017 (Kan/17, H3N2), A/Singapore-IFNIMH-16-0019/2016 (Sing/16, H3N2), A/Hong Kong/4801/2014 (HK/14, H3N2), A/Switzerland/9715293/2013 (Switz/13, H3N2). The H5 virus panel includes: A/Astrakhan/3212/2020 (Ast/20, H5N8) (Virus-like particles, see reference (20) for details), A/Sichuan/26221/2014 (Sich/14, H5N6), A/Guizhou/1/2013 (GZ/13, H5N1), A/Hubei/1/2010 (HB/10, H5N1), A/Egypt/321/2007 (Egy/07, H5N1), A/Whooper swan/Mongolia/244/2005 (Mon/05, H5N1), A/Vietnam/1203/2004 (Viet/04, H5N1).

At 3 weeks after final vaccination, the DBA/2J mice were challenged intranasally with a total volume of 50 μL containing either A/Brisbane/02/2018(H1N1) virus at 3.64×10^4^ PFU or A/Sichuan/26221/2014(H5N6) x PR8 virus at 1×10^6^ PFU. Mice were monitored, at minimum, daily for weight loss, disease signs, and death for 14 days post-inoculation (dpi). Any animal exceeding 25% weight loss or a humane-endpoint score equal to or greater than three was humanely euthanized. Lung samples were harvested on day 3 post-infection (Fig. 1B).

### Hemagglutination-Inhibition (HAI) assay

The HAI assay was used to characterize the antibody response in mouse sera against HA using either 0.8% TRBCs, 0.75% GPRBCs, or 1% HoRBCs according to the WHO laboratory influenza surveillance manual (48). Briefly, the mouse sera was treated with receptor-destroying enzyme (RDE) (Denka Seiken, Co., Japan) at 1:3 ratio at 37°C for 18 h, heat-inactivated at 56°C for 45 min and diluted 1:10 in PBS. The RDE-treated sera was then diluted in a series of two-fold serial dilutions in v-bottom microtiter plates. An equal volume of each virus, adjusted to approximately 8 hemagglutination units (HAU)/50 μL in the presence of 20nM oseltamivir carboxylate (for GPRBCs), was added to each well. The plates were covered and incubated at room temperature for 30 mins, and then red blood cells in PBS were added. The plates were mixed by gentle agitation, covered, and the red blood cells were allowed to settle for 1 h at room temperature. The HAI titer was determined by the reciprocal dilution of the last well that contained non-agglutinated GRBCs.

### Influenza viral plaque assay

MDCK cells were seeded into a six-well plate at 1 × 10^6^ cells/well one day prior to performing the plaque assay. On the day of the assay, frozen lung tissues were thawed on ice, weighed and homogenized in 2 ml of Dulbecco’s modified Eagle’s medium (DMEM) (Thermo Fisher, Waltham, MA, USA). The homogenate was centrifuged at 500 g for 5 min to remove tissue debris, and the supernatant was collected and subjected a serial 10-fold dilution in DMEM supplemented with 1% penicillin-streptomycin (DMEM + P/S) (Thermo Fisher). When MDCK cells reached 90% confluency in each well, the plates were washed 2 times with DMEM + P/S, and infected with 100 µL of each dilution of homogenate supernatant. The plates were then shaken every 15 mins for 1 hour, the supernatant was removed, and cells were washed twice with fresh DMEM + P/S. Following the second wash, a solution of 2 times MEM and 1.6% agarose (Thermo Fisher) mixed 50:50 v/v, and supplemented with 1µg/mL of L-1-tosylamido-2-phenylethyl chloromethyl ketone (TPCK)-treated trypsin (Thermo Fisher) was added into each well. Plates were then incubated at 37°C + 5% CO2 for 48-72 hrs. At the end of incubation, the gel overlays were removed from each well, and the cells were fixed with 10% buffered formalin for 15 mins and stained with 1% crystal violet (Thermo Fisher) for 15 mins at room temperature. Plates were then rinsed thoroughly 3 times with fresh water to remove excess crystal violet. Plates were allowed to air dry for 24 h, and the viral plaques were enumerated as the reciprocal of each dilution. The lung viral titers were calculated and presented as plaque forming units (PFU)/g of lung tissue.

### ELISA for elicited antibody quantification

A high-affinity, 96-well flat-bottom enzyme-linked immunosorbent assay (ELISA; Immulon 4HBX) plate was coated with 100 μL of 1 μg/mL of wildtype rHA in ELISA carbonate buffer (50 mM carbonate buffer [pH 9.5] with 5 mg/mL bovine serum albumin [BSA]), and the plate was incubated overnight at 4°C. The next morning, non-specific epitopes were blocked with 1% BSA in PBS with 0.05% Tween 20 (PBST plus BSA) solution for 1h at RT or overnight at 4°C. Buffer was removed, 3-fold serial dilutions of raw sera were added to the plate with an initial dilution of 1:500 and plates were incubated at 37°C for 90 min. Plates were washed in PBS, goat anti-mouse IgG-HRP (horseradish peroxidase) (cat. no. 1030-05, Southern Biotech, Birmingham, AL, USA) was added at 1:4,000 in PBST plus BSA, and incubated at 37°C for 90 min. After washing, 2,29-azino-bis (3-ethylbenzothiazoline-6-sulfonic acid) (ABTS) substrate in McIlvain’s buffer (pH 5) was added to each well, and incubated at 37°C for 10 min. The colorimetric reaction was stopped with the addition of 1% SDS in dH2O, and the absorbance was measured at 414 nm using a spectrophotometer (PowerWave XS; BioTek, Winooski, VT, USA).

### Statistical analysis

Data is presented as absolute mean values ± standard error of the mean (SEM). The “One-way ANOVA” was used to analyze the statistical differences between groups using GraphPad Prism 9 software (GraphPad, San Diego, CA, USA). A “p” value less than 0.05 was defined as statistically significant (*, P < 0.05; **, P < 0.01; ***, P < 0.001; ****, P < 0.0001).

## Results

### cGAMP MPs adjuvanted COBRA vaccines elicited strong anti-hemagglutinin antibodies

For this study, mice were vaccinated with H1, H3, and/or H5 COBRA HA proteins that were formulated with cGAMP MPs individually or in bivalent or trivalent mixtures (Fig. 1B). Mice vaccinated with the trivalent protein antigen plus cGAMP MPs had high IgG titers against HA proteins from Bris/18 (H1) (Fig. 2A), Tas/20 (H3) (Fig. 2B), and Sich/14 (H5) (Fig. 2C). Similar anti-HA titers were elicited using these same COBRA HA proteins formulated with MF59 pre-clinical mimic AddaVax (Fig. 2). There were no detectable anti-HA antibody titers in mock vaccinated mice and lower anti-HA antibody titers in mice vaccinated with only the COBRA HA antigens without adjuvant or with blank MPs. All mice vaccinated with COBRA HA antigens had primarily anti-HA-specific IgG1 antibodies with lower levels of IgG2a antibodies (Fig. 3A-C). COBRA HA antigen formulated with cGAMP MPs elicited higher ratio of IgG2a antibodies than the COBRA HA antigens without adjuvant or with blank MPs (Fig. 3D). Mice vaccinated with COBRA HA plus AddaVax also had lower anti-HA IgG2a antibody levels than vaccines formulated with cGAMP MPs (Fig. 3A-D).

**Figure 2.**
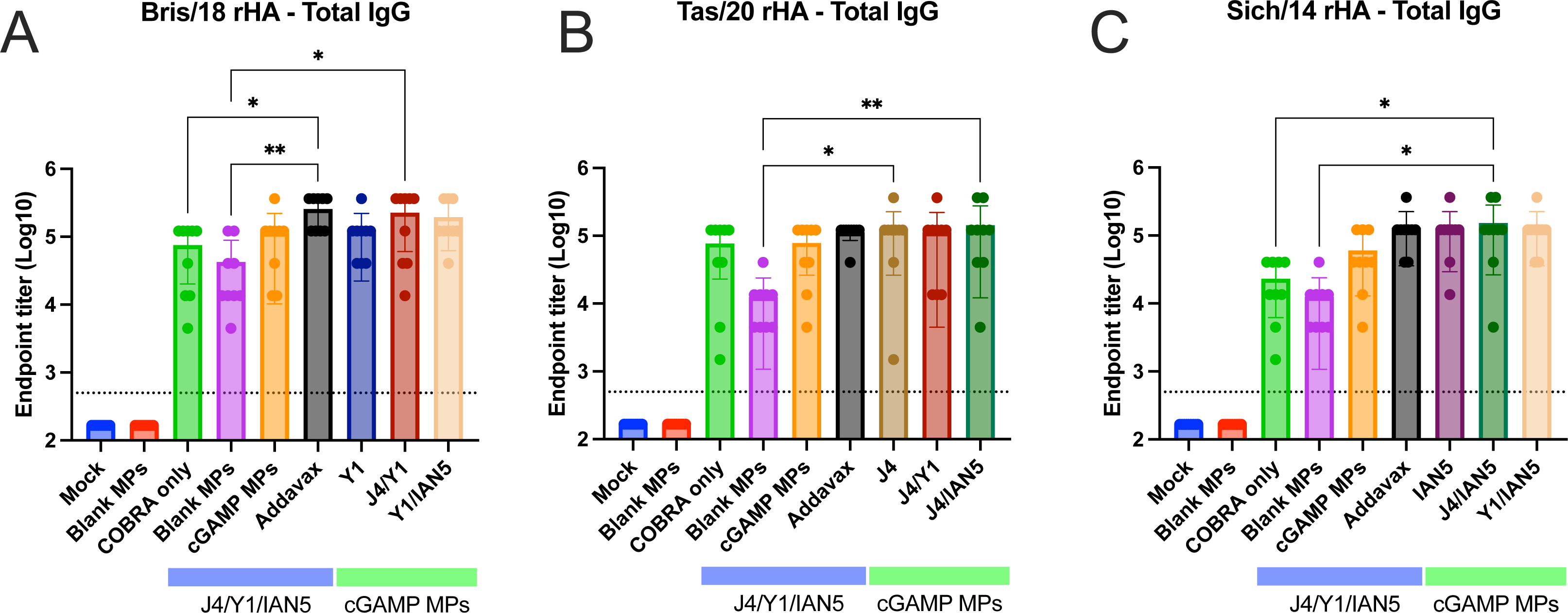
Total IgG antibody response after vaccination. Sera samples collected 2 weeks after final boost were used in ELISA to determine antibody responses against each strain specific HA after vaccination. The following antigens were used: (A) Bris/18 rHA; (B) Tas/20 rHA; and (C) Sich/14 rHA. ELISA data were statistically analyzed using nonparametric one-way analysis of variance (ANOVA) by Prism 9 software (GraphPad Software, Inc., San Diego, CA). A P value of less than 0.05 was defined as statistically significant (*, P < 0.05; **, P < 0.01). Data is presented as average ± standard deviation. The dished line on the graph indicates limit of detection as 1:500.

**Figure 3.**
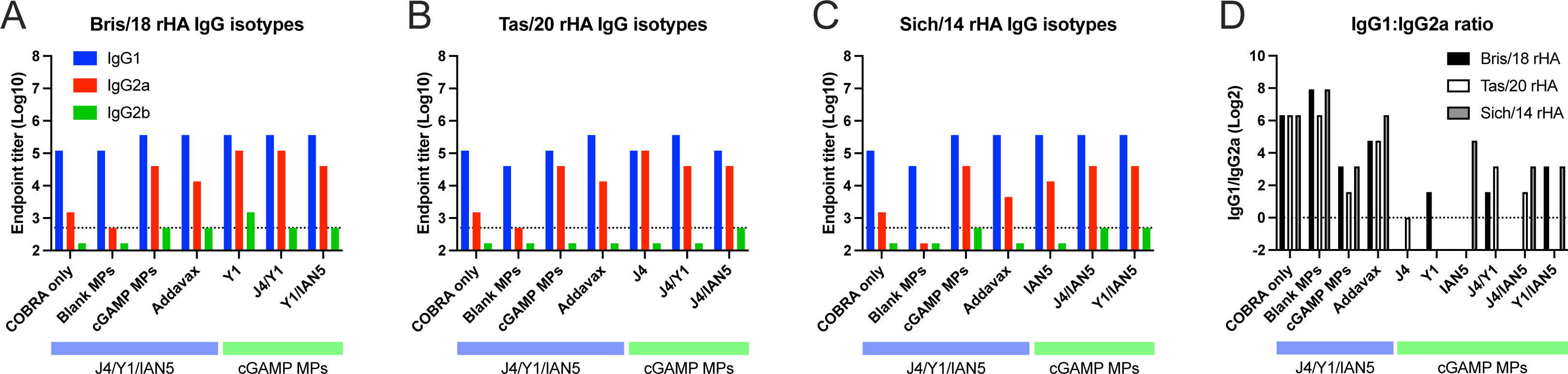
IgG isotype antibody response after vaccination. Sera samples collected 2 weeks after final boost were used in ELISA to determine antibody responses after vaccination. The following antigens were used: (A) Bris/18 rHA; (B) Tas/20 rHA; and (C) Sich/14 rHA. (D) IgG1: IgG2a ratio. The dished line on graphs A, B, and C indicates limit of detection as 1:500. The dished line on graph D indicates IgG1: IgG2a at 1:1 ratio.

Mice vaccinated with the COBRA HA antigens and adjuvanted with cGAMP MPs had sera with HAI activity against the panel of recently isolated H1N1 influenza viruses, but not historical, seasonal H1N1 influenza viruses (SFig. 2). Mice vaccinated with either the trivalent COBRA HA antigens alone or antigens with blank MPs had, on average, HAI titers between 1:40-1:80 against H1N1 influenza viruses (Fig. 4A-D; green and purple bars), but <1:40 against the panel of H3N2 and H5N1 viruses (Fig. 4E-K). Mice vaccinated with these same vaccines plus cGAMP MPs had robust HAI activity against clade 1 and clade 3C.2a H3N2 influenza viruses, but not the viruses in clade 3C.3a where the sera from mice vaccinated with this group displayed low to undetectable HAI activity (SFig. 3). A few mice of all those vaccinated in the experimental group had high HAI titers against the H5Nx viruses of clade 2.3.4.4 and 2.2 (Fig. 4I-K and SFig. 4). Mice vaccinated with the monovalent versions of each COBRA HA antigen with cGAMP MPs had statistically similar HAI titers against each influenza virus as trivalent COBRA HA with cGAMP MPs except mice vaccinated with the monovalent COBRA Y1 (H1) HA antigen with cGAMP MPs. In the COBRA H1 with cGAMP MPs vaccinated mice, significantly higher HAI activity was observed compared to the HAI activity in sera collected from mice vaccinated with the trivalent COBRA HA antigen against Vic/19 and Cal/09 H1N1 influenza viruses (Fig. 4A and D).

**Figure 4.**
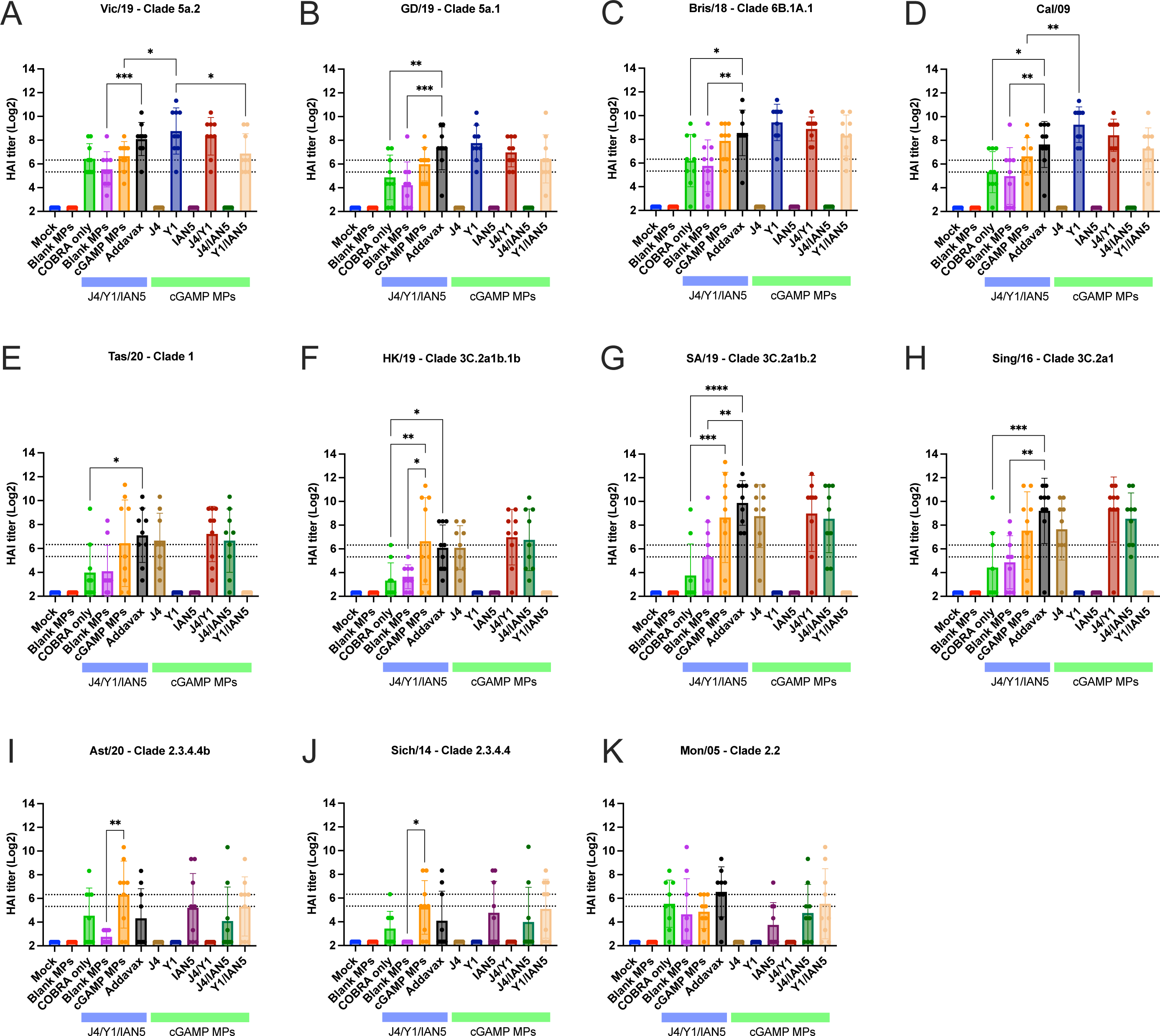
Hemagglutinin inhibition assays. Individual mice serum collected after final boost were used in HAI assay against a panel of historical H1N1, H3N2, and H5Nx influenza viruses. The title of each figure indicates the virus name. The x-axis indicates the experimental group. The y-axis indicates HAI titer in Log_2_. The lower dashed line indicates 1:40 and the higher dashed line indicates 1:80. HAI titers were statistically analyzed using nonparametric one-way analysis of variance (ANOVA). A P value of less than 0.05 was defined as statistically significant (*, P < 0.05; **, P < 0.01; ***, P < 0.001; ****, P < 0.0001). Data is presented as average ± standard deviation.

Mice vaccinated with trivalent COBRA HA antigen with cGAMP MPs or blank MPs had the highest neutralization titers against the Tas/20 and HK/19 H3N2 influenza viruses (Fig. 5A-B and E-F). Similar neutralization titers were observed in trivalent COBRA HA with AddaVax vaccinated mice. In addition, these mice had similar neutralization titers compared to responses from mice vaccinated with COBRA HA proteins with cGAMP MPs in bivalent and monovalent formulations. In contrast, mice vaccinated with unadjuvanted trivalent COBRA HA antigens had lower neutralization log_2_ titers (∼7.5). Vaccine mixtures that lacked the H3 COBRA HA component (J4) had little or no neutralizing titers against two H3N2 influenza viruses tested. Mice vaccinated with any COBRA mixture including the Y1 HA component with or without adjuvant had high neutralization antibody titers against the Bris/18 (H1N1) with a log_2_ titer between 12.5-17.42 (Fig. 5C and G). Similar results were observed against the Sich/14 (H5N6), albeit at lower neutralizing titers, with any mixture including the H5 COBRA component, IAN5 (Fig. 5D and H).

**Figure 5.**
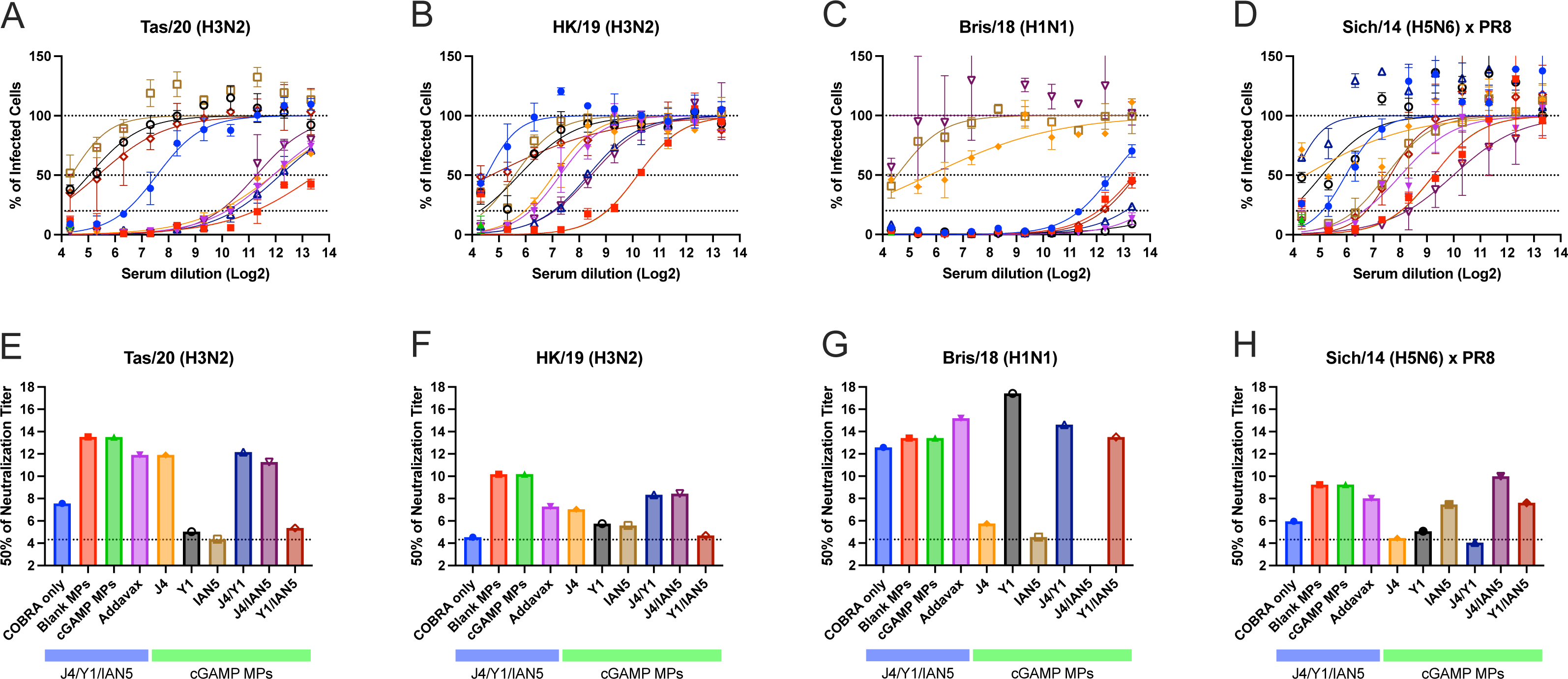
Focal reduction assay (FRA). Pooled mice serum for each group collected after final boost were used in FRA against H3N2, H1N1, and H5N6 influenza viruses. The title of each figure indicates virus name. (A-D) Raw data obtained in FRA. The x-axis indicates the serum dilution in Log_2_, the y-axis indicates the percentage of infected cells by the virus. The lower dotted line represents 80% neutralization (Neut_80_), the middle-dotted line represents 50% neutralization (Neut_50_), and the upper dotted line represents no neutralization of viral infection. Data is presented as average ± standard deviation. (E-H) Neut_50_ titers against each of the indicated viruses. The x-axis indicates the experimental group, the y-axis indicates titer at 50% inhibition in Log_2_. The dished line on the graph indicates limit of detection as 1:20.

### cGAMP MPs adjuvanted COBRA HA antigens protected against lethal influenza challenge

Following Bris/18 (H1N1) virus infection, mock vaccinated mice or mice vaccinated with blank MPs lost more than 20% of their original bodyweight (Fig. 6A), had induction of severe disease (Fig. 6B), and reached the humane endpoint by day 4 to 6 post-infection (Fig. 6C). Mice vaccinated with any formulation containing the H1 COBRA HA protein (Y1) with or without the adjuvant had less than 5% loss of their original body weight, no observed clinical symptoms and survived infection (Fig. 6A-C and SFig. 5A-C). Mice vaccinated with COBRA antigens lacking the Y1 component and challenged with Bris/18 H1N1 lost significant weight, had severe morbidity (SFig. 5A-C), and had high viral lung titers (∼5x10^7^ PFU/g of lung tissue) 3 days post-infection regardless of the inclusion of cGAMP MPs or not (Fig. 6D and SFig. 5D).

**Figure 6.**
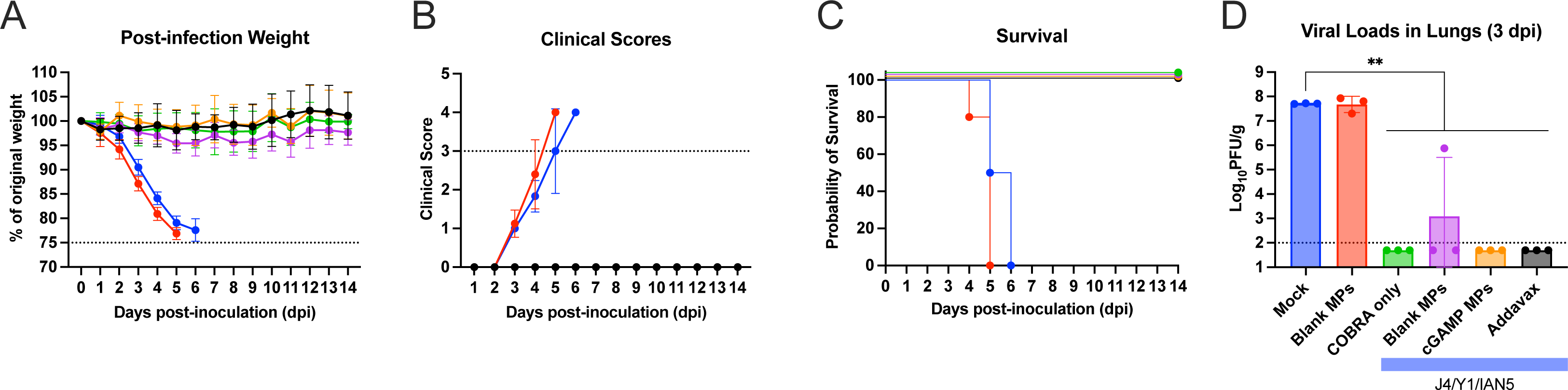
Mice challenged with Bris/18 H1N1 influenza virus. (A) The weight loss curves, (B) survival, (C) clinical scores, and (D) pulmonary viral loads on day 3 post-infection. Colors indicate experimental groups given in D. Data is given as average ± standard deviation. Statistical analysis was conducted using nonparametric one-way analysis of variance (ANOVA). A P value of less than 0.05 was defined as statistically significant (**, P < 0.01).

Mock vaccinated mice infected with Sich/14 (H5N6) virus lost significant bodyweight and all mice reached endpoint by day 6 post-infection (Fig. 7A-C). In contrast, mice vaccinated with trivalent COBRA HA antigens with cGAMP MPs had minimum weight loss (<5%), no clinical symptoms over the course of infection, and all animals survived challenge (Fig. 7A-C). Mice vaccinated with trivalent COBRA HA antigens with blank MPs lost ∼15% on average, and had mild symptoms by day 7 post-infection, but all animals survived (Fig.7A-C). Mock vaccinated mice had the high level of virus present in their lungs (∼5 x 10^6^ PFU/g of lung tissue) (Fig. 7D), which was significantly higher than animals vaccinated with trivalent COBRA HA antigens with cGAMP MPs. There was little or no detectable virus present in the lungs of the mice vaccinated with trivalent COBRA HA antigens with cGAMP MPs. In contrast, mice vaccinated with trivalent COBRA vaccines with blank MPs had lower viral titers in the lung tissue (∼3 x 10^4^ PFU/g of lung tissue) compared to mice in the mock vaccinated group. Animals infected with Tas/20 (H3N2) influenza virus had no weight loss or symptoms observed with all animals surviving challenge and no detectable virus present in their lung tissues.

**Figure 7.**
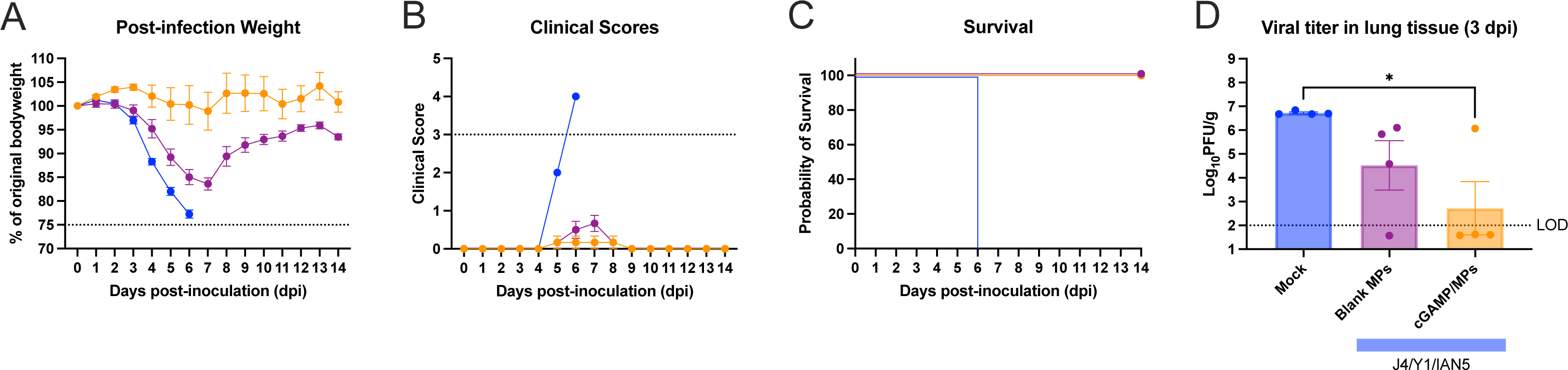
Mice challenged with Sich/14 H5N6 influenza virus. (A) The weight loss curves, (B) survival, (C) clinical scores, and (D) pulmonary viral loads on day 3 post-infection. Colors indicate experimental groups given in D. Data is given as average ± standard deviation. Statistical analysis was conducted using nonparametric one-way analysis of variance (ANOVA). A P value of less than 0.05 was defined as statistically significant (*, P < 0.05).

## Discussion

The goal of many universal influenza vaccines is to elicit immune responses that protect against influenza A viruses in group I and group II (7). To achieve this goal, approaches that direct immune response to conserved protective epitopes on HA, NA, NP, and M2 proteins are under evaluation (8–13). The COBRA design is focused on directing the immune response towards epitopes on the globular head domain of both HA and NA glycoproteins. HA protein sequences from historical and circulating influenza viral isolates were used to generate a COBRA HA protein through a multi-layer consensus building approach (9, 14–16), thus, retaining conserved broadly reactive epitopes (15). The immunity elicited by COBRA hemagglutinins elicit immune responses with a broader coverage of protection compared to wild type HA vaccines (10, 14–21) and the elicited antibodies recognize up to 20 years of future variants (22) across multiple clades (23). In this study, we observed that Y1 COBRA vaccine elicited robust antibody responses against 2009 pandemic-like viruses in a 10-year range, while J4 and IAN5 COBRA HA proteins induced protective antibodies against influenza isolates from multiple clades. These results indicate that the COBRA methodology is an effective vaccine design strategy for universal influenza vaccine development.

Conventional influenza vaccines induce strain-specific immunity and therefore provide limited protection against antigenic drifted strains. To overcome the strain diversity, COBRA-based H1, H3, and H5 HA antigens were generated and evaluated for the breadth of antibody response and protection against influenza viral infection. To ensure potent activation of innate immunity, STING agonist cGAMP formulated in Ace-DEX MPs was co-delivered with COBRA HA proteins and evaluated as a trivalent (H1, H3, H5) influenza vaccine. This study shows that a trivalent COBRA HA vaccine adjuvanted with cGAMP MPs elicited robust HAI responses. A robust HAI response indicates that this vaccine formulation forms antibodies that prevent the attachment of HA to sialic acid receptors on the cell surface, which can therefore lead to a sterilizing viral infection. Trivalent COBRA antigens adjuvanted with blank MPs elicited medium levels of total IgG binding antibodies against H1, H3, and H5 HA antigens. These antibodies had robust HAI activity against H1 viruses, but not H3 and H5 viruses, whereas the antibodies displayed similar capability to neutralize H1, H3, and H5 viruses. This would indicate that COBRA HA proteins delivered with blank MPs can generate a non-sterilizing response, but that antibodies confer protection through other mechanisms, such as antibody-dependent cellular cytotoxicity (ADCC) and complement dependent cytotoxicity (49–51). The total IgG titers and HAI antibodies elicited by vaccination are similar between trivalent COBRA HA antigens alone or with blank MPs. Taken together, these results indicate that the COBRA HA antigens are highly immunogenic and although adjuvant is not required to induce protection, a more robust generation of antibodies and a greater potential for sterilizing immunity is afforded with the inclusion of cGAMP MPs as an adjuvant.

cGAMP MPs rely on STING activation of IRF3 and NF-κB, to then promote the production of type I IFNs and other pro-inflammatory cytokines, driving the activation of innate immune cells and priming of adaptive immune responses. The current study focuses on humoral immune responses and not cellular immune responses after vaccination. It is clear that co-administration of COBRA HA antigens and cGAMP MPs resulted in significantly enhanced antibody titers, but in particular it induced higher levels of Th1 skewing IgG2a compared to AddaVax. Emulsion adjuvants (*e.g.,* MF59 mimic Addavax) are effective inducers of potent Th2-biased responses and humoral immunity, but alone, they fail to induce significant Th1-responses that drive protective immunity against the influenza virus (52). For example, MF59 increases anti-HA neutralizing antibody responses (53), but this increase does not correlate with better protection against influenza viruses (54). This suggests that a Th1 skewed response, which are not effectively activated by squalene emulsions, are necessary for protection (53). Although cellular responses were not specifically examined in this study, cGAMP MPs help to expand germinal center B cells and memory T cells having the potential to induce long-lasting immune responses against COBRA vaccines (34). Furthermore, cellular immune responses afforded by cGAMP MPs are maintained across genetically diverse mice and in mice with immunocompromising conditions, such as obesity (43).

Overall, the inclusion of cGAMP MPs with trivalent COBRA HA proteins during vaccination resulted in the generation of antibodies that were both broadly inhibitory and neutralizing and protected mice against a lethal challenge. COBRA HA proteins formulated with cGAMP MPs elicited robust protective antibody responses against panels of H1, H3, and H5 subtypes of influenza viruses. Antibodies elicited by cGAMP MPs adjuvanted COBRA vaccines mitigated clinical symptoms and decreased pulmonary virus titers in mice challenged with H1 or H5 influenza viruses. Multivalent COBRA HA proteins adjuvanted with cGAMP MPs are a potent influenza vaccine candidate that induces broadly protective immune responses against seasonal and pre-pandemic influenza strains.

However promising trivalent COBRA antigens with cGAMP MPs are as a vaccine formulation, future work will focus on more clinically representative models and enhanced antigen design to advance this platform. Humans are exposed to influenza viruses many times over a lifetime either via infection or vaccination that result in a complex immune history to influenza. The first influenza infection encountered in childhood has long lasting influence on sequential influenza exposures (55) and protection against severe disease from novel HA subtypes in the same phylogenetic group (56). This study was performed in mice that were naïve to influenza, however, our group has published data demonstrating how pre-existing influenza immune imprinting shapes the immune responses to COBRA HA vaccination (16, 23). Further, to expand the protection afforded with our formulation another potential target, such as neuraminidase, can be included. HAI antibodies prevent virus binding, while NA-inhibiting (NAI) antibodies limit release of newly produced viruses, providing another weapon in the vaccine’s protective arsenal. Compared to HA, NA usually has a lower rate of antigenic drift, thus less antigenicity diversity (57, 58). Correlates of protection against heterologous strains have been well documented, potentially leading to a broader acting antigen (59–61). Conserved protective epitopes were also discovered with monoclonal antibodies (62–64). COBRA NA vaccines have broader immune responses than wild type NA vaccines (10, 21) so therefore inclusion of COBRA NA vaccines into the HA formulations may elicit even broader, and more protective immune responses. Future work will not only focus on non-naïve models of influenza infection, but also inclusion of COBRA NA antigens with cGAMP MPs and other adjuvant systems.

## Acknowledgements

We would like to thank James Allen and Ivette Nuñez for designing COBRA HA sequences and Spencer Pierce for purifying the HA proteins. We thank Naoko Uno for technical assistance. We also thank the University of Georgia Animal Resource staff, technicians, and veterinarians for the excellent animal care. This work was performed in part at the Chapel Hill Analytical and Nanofabrication Laboratory, CHANL, a member of the North Carolina Research Triangle Nanotechnology Network, RTNN, which is supported by the National Science Foundation, Grant ECCS-2025064, as part of the National Nanotechnology Coordinated Infrastructure, NNCI. This project has been funded as part of the Collaborative Influenza Vaccine Innovations Centers (CIVICs) by the National Institute of Allergy and Infectious Diseases, a component of the NIH, Department of Health and Human Services, under contract 75N93019C00052. TMR is also supported in part as a Georgia Eminent Scholar by the Georgia Research Alliance, GRA-001. The work is also supported by R01AI147497 (Ainslie).

## Author Contributions

Conceptualization, X.Z., D.A.H., E.M.B., K.M.A., and T.M.R.; formal analysis, X.Z., H.S. and T.M.R.; investigation, X.Z. and H.S.; resources, K.M.A., T.M.R.; data curation, X.Z.; writing— original draft preparation, X.Z.; writing—review and editing, X.Z., D.A.H., K.M.A. and T.M.R.; supervision, T.M.R.; project administration, X.Z. and H.S.; funding acquisition, K.M.A., T.M.R. All authors have read and agreed to the published version of the manuscript.

## Competing Interests

The authors declare no competing interests.

## Data Availability

The data that support the findings of this study are available from the corresponding author upon reasonable request.

---

**Supplementary Figure 1.** Scanning electron micrographs of (A) blank Ace-DEX MPs and (B) cGAMP loaded Ace-DEX MPs prepared via the electrospray method. Scale bars represent 5 µm.

**Supplementary Figure 2. Hemagglutinin inhibition assays H1 virus panel.** Pooled mice serum collected after each vaccination were used in HAI assay against a panel of historical H1N1 influenza viruses. The title of each figure indicates the virus name. The x-axis indicates the experimental group. The y-axis indicates HAI titer in Log_2_. The lower dashed line indicates 1:40 and the higher dashed line indicates 1:80.

**Supplementary Figure 3. Hemagglutinin inhibition assays H3 virus panel.** Pooled mice serum collected after each vaccination were used in HAI assay against a panel of historical H3N2 influenza viruses. The title of each figure indicates the virus name. The x-axis indicates the experimental group. The y-axis indicates HAI titer in Log_2_. The lower dashed line indicates 1:40 and the higher dashed line indicates 1:80.

**Supplementary Figure 4. Hemagglutinin inhibition assays H5 virus panel.** Pooled mice serum collected after each vaccination were used in HAI assay against a panel of historical H5Nx influenza viruses. The title of each figure indicates the virus name. The x-axis indicates the experimental group. The y-axis indicates HAI titer in Log_2_. The lower dashed line indicates 1:40 and the higher dashed line indicates 1:80.

**Supplementary Figure 5. Mice challenged with Bris/18 H1N1 influenza virus. .** (A) The weight loss curves, (B) survival, (C) clinical scores, and (D) pulmonary viral loads on day 3 post-infection. Colors indicate experimental groups given in D. Data is given as average ± standard deviation. Statistical analysis was conducted using nonparametric one-way analysis of variance (ANOVA). A P value of less than 0.05 was defined as statistically significant (*, P < 0.05; **, P < 0.01; ***, P < 0.001; ****, P < 0.0001).

